# Impact of juglone on oral carcinogenesis induced by 4-nitroquinoline-1-oxide (4NQO) in rat model

**DOI:** 10.1101/2024.01.26.577505

**Authors:** Olgun Topal, Burcu Topal, Yunus Baş, Bünyamin Ongan, Gökhan Sadi, Esra Aslan, Betül Demirciler Yavaş, Mehmet Bilgehan Pektaş

## Abstract

In this study, the potential effects of juglone, also known as PIN1 inhibitor, on oral cancer and carcinogenesis were investigated at the molecular level. 4-Nitroquinoline N-oxide (4-NQO) was used to create an oral cancer model on animals. *Wistar* rats were divided into five groups; Control, NQO, Juglone, NQO+J, and NQO+J*. The tongue tissues of the rats were isolated after the experiment, morphological changes were investigated by histological examinations, and the molecular apoptotic process was investigated by *rt-qPCR* and *Western blot*. Histological results indicate that tumors are formed in the tongue tissue with 4-NQO, and juglone treatment largely corrects the epithelial changes that developed with 4-NQO. It has been determined that apoptotic factors p53, Bax, and caspases are induced by the effect of juglone, while antiapoptotic factors such as Bcl-2 are suppressed. However, it was observed that the positive effects were more pronounced in rats given juglone together with 4-NQO. The use of PIN1 inhibitors such as juglone in place of existing therapeutic approaches might be a promising and novel approach to the preservation and treatment of oral cancer and carcinogenesis. However, further research is required to investigate the practical application of such inhibitors.

## Introduction

Oral cancer is the most common type of cancer in the head-neck region, and recent data predict that its incidence will increase in the coming years [1]. In particular, smoking and consuming alcohol are the main factors contributing to the risk of oral cancer [2]. In terms of clinical pathogenesis, poor oral hygiene, loose dentures, chronic irritation of teeth, nutritional deficiencies, ultraviolet light, radiation exposure, viral infections including *Human papillomavirus* (HPV), *Epstein-barr* virus (EBV), *Herpes simplex* virus (HSV), and fungal infections, including *Candida albicans*, are known to cause oral carcinogenesis [3]. Oral carcinogenesis involves a series of complex simultaneous stages that include the development of precancerous lesions and the processes of invasion and metastasis [4]. The main challenges in oral cancer can be listed as late diagnosis and drug resistance [5,6]. Head and neck cancers is a term that includes the oral and nasal cavities, pharynx, larynx, and paranasal sinuses; however, there is no standard definition of the term oral cancer, as there is no clear distinction regarding whether some anatomical regions (such as the base of the tongue and soft palate) are included in the oral cavity or oropharynx. The oral cavity generally includes the mucosal part of the lip, the anterior 2/3 of the tongue, the floor of the mouth, the gingiva, the retromolar triangle, the buccal mucosa and the hard palate [7]. Squamous cell carcinomas constitute 90% of oral cancers, and the five-year survival rate in this type has been reported to be approximately 56% [8]. It is known that many oral squamous cell carcinomas develop in association with a precancerous lesion, especially leukoplakia, or when these lesions undergo malignant transformation. It has been reported that it most commonly develops in the posterolateral or ventral tongue regions of the mouth. Other areas where it is seen are the floor of the mouth, soft palate, gingiva, cheek, lip, and hard palate [9]. The degree of the disease and the location of the tumor are important in the treatment planning of oral squamous cell carcinoma. Primary treatment is extensive surgical resection. Surgery may be followed by radiotherapy and/or chemotherapy depending on the pathological character of the tumor. While radiotherapy is not thought to be beneficial in early stage carcinomas (stage I and II) with negative surgical margins; III - IV. Tumors with stage, positive surgical margins, or lymph node involvement with one or more extra capsular spread are considered to have a high risk of recurrence, and radiotherapy or chemo-radiation is recommended following surgery [10]. Chemotherapy can usually be applied together with radiotherapy, as initial treatment before chemo-radiation, or as palliative treatment. In addition, in the last 30 years, there have been treatment approaches in which only chemotherapy and radiotherapy are applied for ‘organ preservation’ purposes, especially in larynx and oropharynx carcinomas. Survival rates of primary chemo-radiotherapy have been reported in a wide range of 29-66% [11]. Due to the potential side effects and morbidity risks of drugs used in the field of cancer, the researches for treatment with natural or herbal products constitutes an important source of studies in the development of chemotherapeutic strategies [12]. In this context, recent studies point to the potential effects of phytotherapeutics that inhibit PIN1 in cancer treatment. Under stress, cells release pro-apoptotic proteins such as Bax (Bcl-2-associated X protein) and caspases to initiate apoptosis under the control of cytosolic p53 (Tumor protein 53). These proteins also deactivate these structures by forming complexes with anti-apoptotic factors such as Bcl-xl (B-cell lymphoma xl) and Bcl-2 (B-cell lymphoma 2). This entire process is catalyzed by the PIN1 enzyme. Thus, apoptosis is controlled through conformational change of cytosolic p53 [13].

Juglone is a naphthoquinone derivative PIN1 inhibitor obtained from the plant *Juglans mandshurica* Maxim. Recent studies point to the anti-inflammatory, cytotoxic, anti-tumor, antibacterial, and antioxidant effects of juglone [14–16]. Importantly, juglone, which is known to induce the apoptotic process through PIN1 inhibition, has been shown to have anti-tumor effects in pancreatic, prostate, melanoma, colorectal, glioblastoma, and breast cancers [17–22]. Although there are many studies on the tumor suppressor effects of juglone, its effects on oral cancer types remain unknown. Therefore, in this study, we aimed to investigate both the prophylactic and therapeutic effects of juglone by creating an oral cancer model with 4-NQO in rats.

## Materials and methods

### Animals

The study protocol was approved by the Ethical Animal Research Committee of Afyon Kocatepe University (49533702/291) and it was designed in accordance with the ARRIVE guidelines/checklist. Three-month-old, male, approximately 300 g, *Wistar* 60 rats were obtained from Afyon Kocatepe University Experimental Animal Application and Research Center. Standart rodent chow (Korkutelim Yem Sanayi A.ş., Antalya/Türkiye) that composed of 62% starch, 23% protein, 4% fat, 7% cellulose, vitamins, and salt mixture were given ad libitum with drinking water during the experiment. Before practical applications, rats were acclimatized for one week, and then five groups were formed as: Control, NQO, Juglone, NQO+J (those given juglone after creating an oral cancer model), and NQO+J*.

### Protocols

4-NQO (4-Nitroquinoline N-oxide) solution was prepared as 50 ppm (0.05 mg/ml) in drinking water. After the health status of the rats was checked, 4-NQO was administered to the NQO, NQO+J, and the NQO+J* groups in the drinking water of rats for 8 weeks. Juglone was dissolved in 100% ethanol 14 mM (2.44 mg/ml stock solution). A dosage adjustment was made to achieve 1 mg/kg body weight in a final volume of 2 ml per injection with saline. Juglone injections were applied to the NQO+J group after being given 4-NQO for 8 weeks, and it was applied to the NQO+J* group together with 4-NQO application for 8 weeks. However, 2 ml saline injections were applied to the Control and NQO groups for 8 weeks. At the end of the experiment, the animals were anesthetized and decapitated with a combination of ketamine (100mg/kg) and xylazine (10mg/kg). Tongue tissue samples from the implant site were isolated from animals and stored at −85°C for rt-PCR measurements; for histological analysis, it was stored in 10% formalin solution.

### Determination of the gene expressions with rt-qPCR

Total RNAs were isolated from 50 mg of tongue tissues with RNeasy total RNA isolation kit (Qiagen, Venlo, Netherland) as described in the manufacturer protocol. After isolation, amount and the quality of total RNA were determined by spectrophotometry at 260/280 nm with ThermoScanGO microplate reader (Thermo Fisher Scientific, USA). Presence of any RNA degradation was checked with agarose gel electrophoresis. cDNA synthesis was carried out by using 500 ng of total RNA and oligo (dT)18 primer using commercial first strand cDNA synthesis kit (Thermo Scientific, USA) as described by the supplier. Gene expressions of interested proteins were determined with real time PCR by mixing 1 μl cDNA, 5μl SYBR Green Mastermix (Roche FastStart Universal, SYBR Green Master Mix) and primer pairs (Table 1) at 0.5 μM final concentrations each in a final volume of 10 μl. The real time PCR program of the quantitative PCR (LightCycler480 II, Roche, Basel, Switzerland) was arranged as follows: initial denaturation at 95°C for 10 min, denaturation at 95°C for 10 s, annealing at 58°C for 15 s and extension at 72°C for 15 s with 40 times repeated thermal cycle measuring the green fluorescence at the end of each extension steps. PCR reactions were carried out in triplicates and specificity of PCR products were checked out by melt analysis. Also, negative controls lacking template were used in all reactions. Relative expressions of genes with respect to internal control GAPDH were calculated with the efficiency corrected advance relative quantification tool of the LightCycler® 480 SW 1.5.1 software.

**Table 1.**
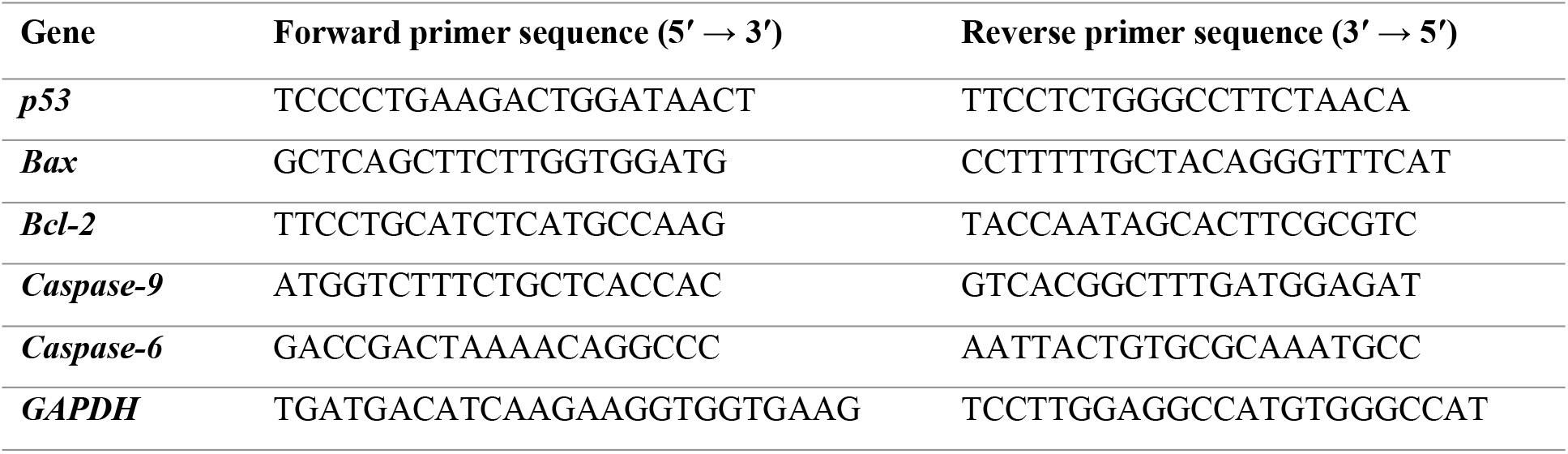
Primer sequences of p53, Bax, Bcl-2, Caspase-9, Caspase-6, and the internal standard GAPDH used for the mRNA expression determination of qRT-PCR.

### Determination of protein expressions by Western blot

For the determination of p53, Bax, Bcl-2, Caspase-9, and Caspase-3 protein contents, tongue tissue was homogenized in 2-fold volumes of homogenization medium (50 mM Tris pH:7.4, 150 mM NaCl, 5 mM EDTA, 1 % (w/w) Triton X-100, 0.26% (w/v) sodium deoxycholate, 50 mM sodium fluoride, 0.1 mM sodium orthovanadate and 0.2 mM phenylmethylsulfonyl fluoride (PMSF)) with a Tissue RuptorTM (Qiagen, Netherlands) homogenizer. The homogenates were centrifuged at 1500 x g for 10 min at 4ºC. After the removal of the supernatants, the protein concentrations were determined by the Lowry method [23]. Microgram of total proteins were separated by SDS-PAGE and transferred on to PVDF membranes using a semi-dry electroblotting apparatus (TransBlot Turbo, BioRad, Germany). Blotted membranes were then blocked with 5% (w/v) nonfat dried milk and incubated with primary antibodies for p53 (Anti-TP53 antibody, 1/1000), Bax (Anti-Bax antibody, 30-110 Internal, 1/1000), Bcl-2 (Anti-Bcl-2 antibody, 40-120, 1/1000), caspase-9 (Anti-CASP9 antibody, 60-140, 1/1000), and caspases-3 (Anti-CASP3 antibody, 1/1000) for 2 h at room temperature or overnight at 4ºC. As an internal control, GAPDH proteins were also labeled with anti-GAPDH Rabbit IgG (Santa Cruz, sc:25778, 1:2,000). Horseradish peroxidase-conjugated secondary antibody (Santa Cruz, sc:2030 or sc:2770, 1:10,000) was incubated for 1 h, and the blots were incubated in Clarity™ Western ECL (Bio-Rad Laboratories, Hercules CA, USA) substrate solution. Images of the blots were obtained using the ChemiDoc™ MP Chemiluminescence detection system (Bio-Rad Laboratories, Hercules CA, USA) equipped with a CCD camera. The relative expression of proteins with respect to GAPDH was calculated using ImageLab5.2 software.

### Histochemical staining

The tongue and tongue roots taken from the rats were fixed in 10% neutral formalin and then histologically processed and embedded longitudinally in paraffin. Then, 5μ sections were taken on classic slides; Hematoxylin-eosin staining was applied all the sections and evaluated under a light microscope.

For histopathological evaluation of the epithelium, the epithelium and cell structure were evaluated and a 6-grade scoring was made based on the WHO classification of oral epithelial dysplasia reported by Speight [24]. The classification model was described in Table 2. Additionally, hyperkeratinization and vascular dilatation formation were examined using the evaluation method of Ross et al [25]. The areas where hyperkeratinization and vasodilation were most intense in the sections were determined and these changes were classified into three grades: mild (1), moderate (2) and severe (3).

**Table 2.** Morphological changes according to Speights’ scores.

| *Morphology* | *Score* | *Cell structure* | *Epithelial structure* |
| --- | --- | --- | --- |
| *Normal epithelium* | 0 | No change | No change |
| *Hyperplasia* | 1 | No change | Although normal maturation, epithelial thickening and hyperkeratosis |
| *Mild dysplasia* | 2 | Differentiation in cell and nucleus shapes in the lower 1/3 of the epithelium | Thickening in the lower 1/3 of the epithelium, disruption of cell orientations |
| *Moderate dysplasia* | 3 | Differentiation in cell and nucleus shapes up to the middle 1/3 of the epithelium | Increase in the number of cells up to the middle 1/3 of the epithelium, thickening of the epithelium, complete disruption of cell orientation |
| *Severe dysplasia* | 4 | Disruption of the full-thickness cell structure of the epithelium | Migration of atypical cells from the basal lamina to the lamina propria in addition to the disrupted epithelial structure |
| *Carcinoma in situ* | 5 | All changes available | Full-layer structural changes and delamination |

**Table 3.** Histopathological changes of the tongue tissues from the Control, NQO, Juglone, NQO+J, and the NQO+J* groups.

| Groups | Oral Epithelial Dysplasia | Hyperkeratinization | Vascular Dilatation |
| --- | --- | --- | --- |
| Control | 0 | 1 | 1 |
| NQO | 3.57±0.57 | 2.83±0.17 | 2.67±0.21 |
| Juglone | 0.84±0.31 | 1.16±0.17 | 1 |
| NQO+J | 1.33±0.21 | 1 | 1.5±0.22 |
| NQO+J* | 1.67±0.67 | 1.5±0.34 | 1.33±0.33 |

### Statistical analysis

Gene and protein expressions of all groups were normalized to the mean of the Control groups and data was also normalized with corresponding GAPDH. All data is represented as mean ± standard error of the mean (SEM) through the study. Statistical comparisons were performed using one-way ANOVA followed by appropriate post-hoc test (Tukey). Comparisons giving *p* values less than 0.05 were accepted as statistically significant.

## Results

### The influences of juglone in the mRNA expressions of the apoptosis

As shown in Fig. 1, the gene expression levels of p53 (a), Bax (b), Bcl-2 (c), caspases-9 (d), and caspases-6 (e) in the tongue tissues from rats were established by *realtime PCR* analysis. 4-NQO-feeding (NQO group) decreased p53, Bax, and caspases-9 and increased Bcl-2 mRNA expressions in the tongue tissue samples from rats, whereas no changes were observed in caspases-6 mRNA expressions (*p* < 0.05 compared to Control). Intraperitoneal juglone injection (Juglone group) significantly increased p53, Bax, caspase-9, and caspases-6 mRNA expressions in the tongue tissues of healthy rats, whereas markedly reduced Bcl-2 expressions (*p* < 0.05 compared to control). Juglone administration together with NQO-feeding (NQO+J* group) significantly increased p53, Bax, caspases-9, and caspases-6 mRNA expressions and decreased Bcl-2 levels in the tongue tissue samples from rats compared to the NQO groups. Similarly, juglone injection after the carcinogenesis induced by 4-NQO (NQO+J group), Bax and caspases-9 mRNA expressions, while decreased Bcl-2 levels but no changes were observed in p53 and caspases-6 expressions in the tongue tissue samples from rats compared to the NQO groups. On the other hand, caspase-9 mRNA expressions in rats given juglone along with 4-NQO (NQO+J* group) increased significantly more than given juglone after carcinogenesis (NQO+J group). No similar significant change was observed in other parameters.

**Figure 1.**
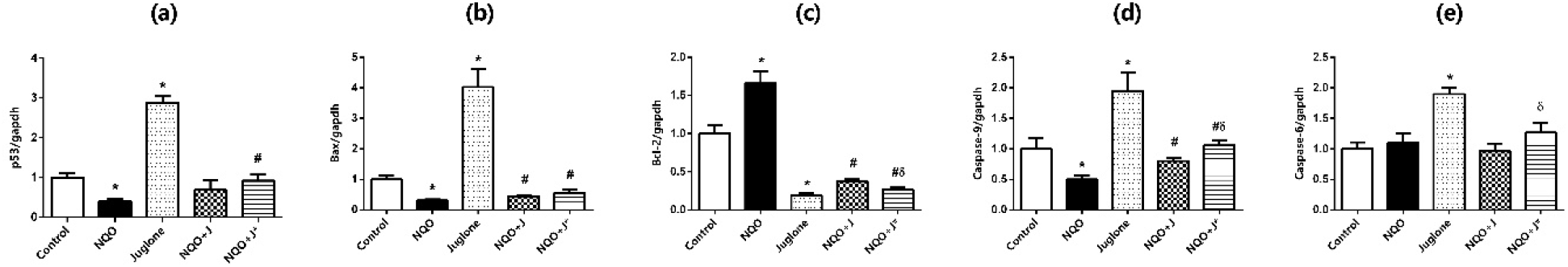
mRNA expression levels of p53 (a), Bax (b), Bcl-2 (c), Caspase-9 (d), and Caspase-6 (e) in the tongue of rats from the Control, NQO, Juglone, NQO+J, and the NQO+J* groups. Data was normalized by GAPDH. Each bar represents the means of at least six rats. Values are expressed as mean ± SEM. * Significantly different (p < 0.05) compared to Control group; # significantly different (p < 0.05) compared to NQO group; δ significantly different (p < 0.05) compared to NQO+J group.

### The influences of juglone in the protein expressions of the apoptosis

As shown in Fig. 2, the protein expression levels of p53 (a), Bax (b), Bcl-2 (c), caspases-9 (d), and caspases-3 (e) in the tongue tissues from rats were established by *Western blot* analysis. 4-NQO-feeding (NQO group) decreased p53, Bax, caspases-9, and caspases-3 levels and increased Bcl-2 protein expressions in the tongue tissue samples from rats (*p* < 0.05 compared to Control). Intraperitoneal juglone injection (Juglone group) significantly increased p53, Bax, and caspases-3 protein levels in the tongue tissues of healthy rats, and markedly reduced Bcl-2 levels, whereas no changes were found in the caspase-9 expressions (*p* < 0.05 compared to Control). Juglone administration together with NQO-feeding (NQO+J* group) markedly increased p53, and caspases-9 levels and significantly decreased Bcl-2 levels, whereas no changes were observed in the Bax and caspase-3 expressions in the tongue tissue samples from rats compared to the NQO groups. Similarly, juglone injection after the carcinogenesis induced by 4-NQO (NQO+J group), caspases-9 levels were found to be increased compared to NQO group, whereas no changes were observed in the other parameters. On the other hand, in rats given juglone along with 4-NQO (NQO+J* group) markedly increased p53 expressions and significantly reduced Bcl-2 levels more than given juglone after carcinogenesis (NQO+J group). No similar significant change was observed in other parameters.

**Figure 2.**
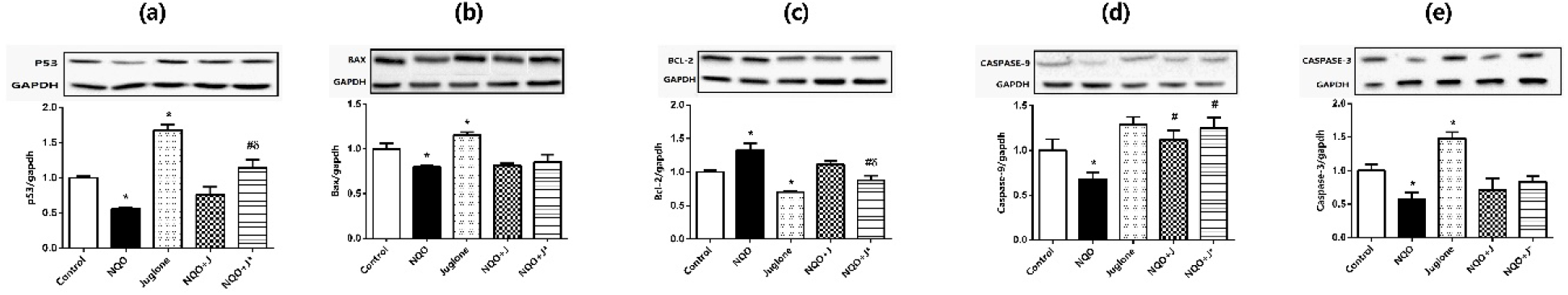
The protein expression levels of insulin receptor substrate p53 (a), Bax (b), Bcl-2 (c), Caspase-9 (d), and Caspase-3 in the tongue of rats from the Control, NQO, Juglone, NQO+J, and the NQO+J* groups. The protein levels were quantified using densitometry and normalized with GAPDH. Representative Western blot images are included above the corresponding figures. Each bar represents at least six rats. * Significantly different (p < 0.05) compared to Control group; # significantly different (p < 0.05) compared to NQO group; δ significantly different (p < 0.05) compared to NQO+J group.

**Figure 3.**
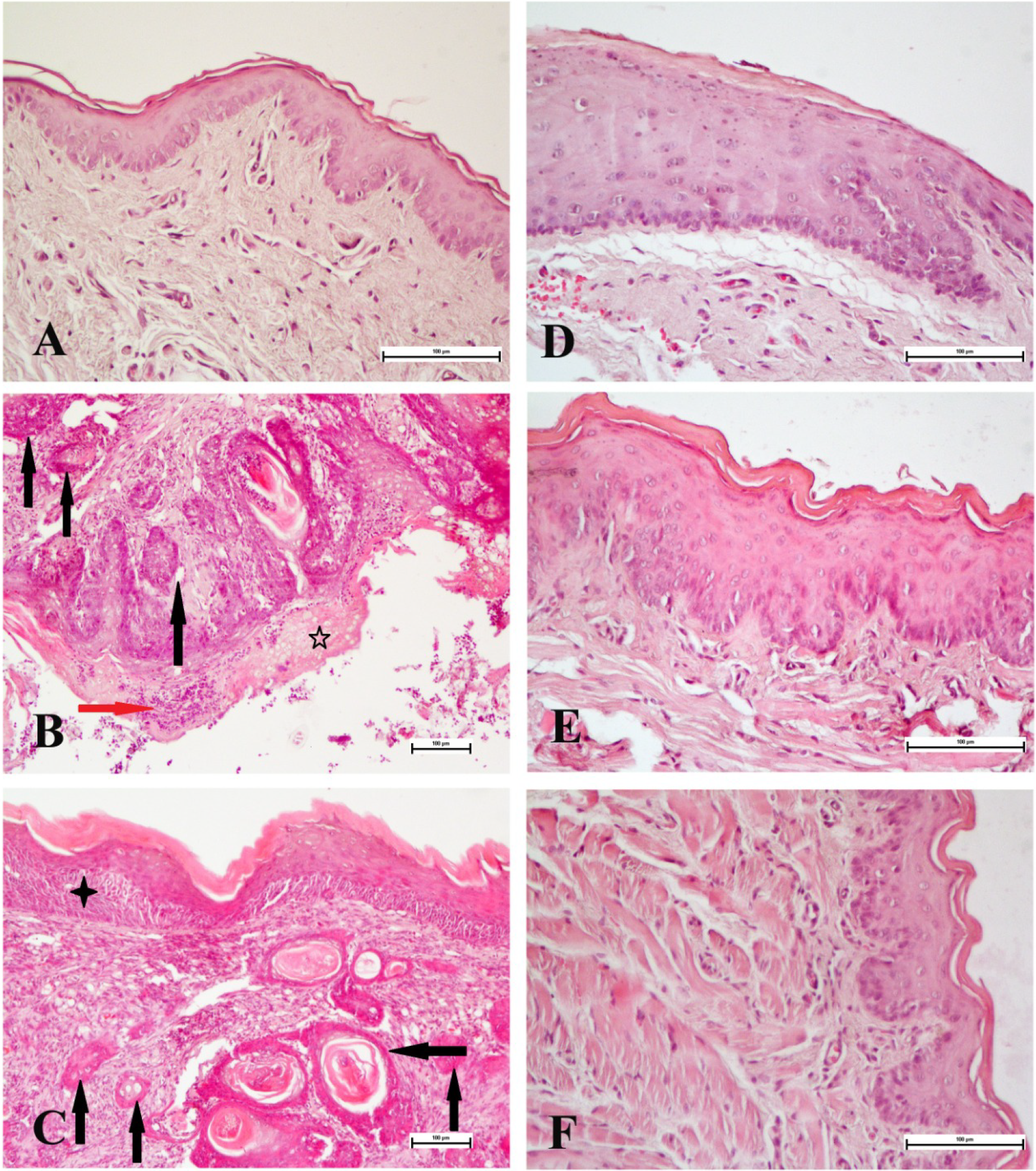
Histopathological features of the tongue tissues from the Control (A), NQO (B and C), Juglone (D), NQO+J (E), and the NQO+J* (F) groups. Hematoxylin-Eosin staining A, D, E, and F x200, B and C x100; Scale bar: 100 μm. While the Control and Juglone groups showed normal morphology, epithelial changes and tumor formation are observed in the NQO group (*Black arrows*: Tumoral areas; *Red arrow*: Lymphocyte infiltration; *Five-pointed star*: hyperkeratinization; *Four-pointed star*: Carcinoma *in situ*). It is observed that epithelial changes are significantly reduced in NQO+J and NQO+J* groups.

### The influences of juglone on histological parameters

As a result of histopathological evaluation under a light microscope, it was observed that the cell arrangements in the Control and Juglone groups were regular and there was no atypia in their nuclei. However, while hyperkeratotic changes and epithelial thickening were observed in some of the tissues in the Juglone group, mild dysplasia was observed in only one sample. Again, hyperkeratinization and vascular dilatation were observed to be mild in these groups. While moderate dysplasia was observed extensively in the NQO group, *carcinoma in situ* was observed to develop in one sample. Again, severe hyperkeratinization and severe vascular dilatation were observed in this group. While hyperplasia was observed in most of the samples in the NQO+J group, carcinoma in situ was observed in one sample. While mild hyperkeratinization was observed in this group, moderate vascular dilatation was observed. In the NQO+J* group, while most of the samples showed hyperplasia, mild dysplasia was observed in one sample. Hyperkeratinization and vascular dilatation were observed to a mild degree.

## Discussion

Oral cancer consists of cancer of areas of the oral cavity including the lips, labial and buccal mucosa, anterior two-thirds of the tongue, maxillary and mandibular gums, retromolar region, floor of the mouth under the tongue, and roof of the mouth. These areas are histologically covered with squamous epithelium (keratinized/non-keratinized, masticatory, special mucosa). The remainder consists of oral cancer, verrucous carcinoma, minor salivary gland malignancies, mucosal melanoma, Kaposi’s sarcoma, primary intraosseous squamous cell carcinoma, osteosarcoma, odontogenic tumors, metastatic tumors, and connective tissue tumors [26]. One of the basic features that the cell often, but not always, acquires is the ability to escape from apoptosis. Apoptosis, technically called programmed cell death, is an important element in the pathogenesis of many diseases [27]. It is known that the initiation of apoptosis occurs via trans-membrane receptor-mediated interactions or caspase-initiated mitochondria. According to Jorge Finnigan, TNF (ligands and receptors), p53 family, Bax/Bcl-2 family, as well as caspases, are noteworthy among the genes involved in apoptosis [28]. Recent studies have focused on the role of PIN1 inhibition in this process. PIN1 is a member of a group of three prolyl isomerases. PIN1 interacts with the motif containing the phospho-Ser/Thr-Pro of substrates and increases the cis-trans isomerization of peptide bonds, thereby controlling the functions of these substrates. Importantly, PIN1 expression level is highly upregulated in most cancer cells and is associated with malignant features and therefore poor outcomes. Additionally, PIN1 was revealed to promote the functions of multiple oncogenes and eliminate tumor suppressors. Accordingly, PIN1 is well known as a key regulator of malignant processes [29]. It is clearly stated that PIN1 activation has an anti-apoptotic effect [30]. It has also been shown to have a role in the regulation of caspases [31]. Hyperactivated PIN1 has been shown to cause nasopharyngeal carcinoma [32]. A study conducted in Eastern China found that the risk of oral squamous cell carcinoma was significantly increased in 209 patients with PIN1 polymorphism [33]. Therefore, it can be said that juglone, a strong PIN1 inhibitor, may be a potential candidate for treatment in oral cancer models. In this study, an oral carcinoma model was tried to be created with 4-NQO, and some parameters in the apoptotic pathway were measured by administering juglone both during this process and after 4-NQO application. According to the study results, it was determined that the apoptotic factors p53, Bax, and caspase-9 mRNA and protein expressions, as well as caspase-3 enzyme levels, were suppressed with 4-NQO, while the anti-apoptotic Bcl-2 mRNA and protein expressions were increased. In addition, histological examinations showed intense dysplasia with 4-NQO, hyperkeratinization, and severe vascular dilatation, in one sample, *carcinoma in situ*. These results indicate that an oral carcinoma model was formed with 4-NQO, in line with the literature [34]. It has been shown that juglone administration causes Bax hyperactivation in the Ehrlich carcinoma model developed on male BALB/c mice [35]. In a similar study, it was shown that juglone fractions caused the induction of p53, caspase-9, caspase-3, and Bax, while suppressing Bcl-2 levels [15]. In the nasopharyngeal carcinoma model created by PIN1 induction, juglone was shown to increase caspase-3 protein expression [36]. In our study, juglone was applied for prophylactic and therapeutic purposes at a dose compatible with the literature (1 mg/kg) [37]. Study results indicate that Bax and caspase-9 mRNA expressions from apoptotic structures increased by administering juglone to healthy rats together with 4-NQO and to rats with oral carcinoma induced by 4-NQO, while p53 levels increased significantly in the 4-NQO+J* group, and there is an increasing trend in the 4-NQO+J group. Additionally, juglone was found to dramatically suppress Bcl-2 mRNA expression in all groups. When the protein expressions are evaluated, it is understood that the results are largely parallel to mRNA expressions, and as a different factor, caspases-3 values are also induced by juglone application. It can be said that our results are compatible with the effects of juglone in other cancer models. Another point worth noting here is that the anti-carcinogenic effects of juglone are more pronounced at the prophylactic level (when given together with 4-NQO). This may be particularly related to the regulatory activity of PIN1 on apoptotic/anti-apoptotic structures. The fact that the effects of juglone are more aggressive at the scope of the mRNAs in healthy rats raises questions about the safety of juglone. In a study on the subject, it was stated that the IC50 values of juglone can kill prokaryotic cells without damaging eukaryotic cells, thus juglone can be used in pharmaceutical fungicidal and bactericidal applications, as well as food safety applications and agriculture [38]. In our study, we can say that although juglone’s apoptosis was clearly expressed at the based on the mRNAs, this was not reflected in protein synthesis to the same extent. This may be related to the fact that apoptosis is regulated through different pathways.

While dysplasia develops histologically in oral carcinoma, hyperkeratinization, severe vascular dilatation, and even *carcinoma in situ* are likely to be observed [34]. In our study, the histological findings obtained by examinations under a light microscope were evaluated with Speights’ scoring [39]. The results showed that low-level hyperkeratotic changes and epithelial thickening occurred with juglone application in healthy rats. It was determined that severe hyperkeratinization, high dysplasia and vascular dilatation that developed with 4-NQO were alleviated in both groups administered juglone. In parallel with the molecular results, these changes were found to be more pronounced in rats given juglone together with 4-NQO.

Current data suggest that the possible effects of juglone may be mediated by regulation of the apoptosis signaling pathway due to PIN1 inhibition. Although there are limited studies on the effects of juglone, our findings were found to be compatible with juglone’s pancreatic, prostate, melanoma, colorectal, glioblastoma, breast, Ehrlich ascites, and nasolaryngeal cancer models [15,17,19–22]. Further studies are needed to substantiate our hypothetical assessments.

## Conclusions

Oral carcinoma has become one of the most common types of cancer in recent times. Chemotherapeutics used in treatment have pushed researchers to alternative methods because they carry the risk of morbidity. Juglone is a natural agent that mediates PIN1 inhibition, offering a different approach in cancer treatment. There are no studies examining the effects of juglone on oral carcinoma. Our study results indicate that juglone provides strong anti-carcinogenic activity on oral carcinoma, as well as on some types of cancer, and shows this activity both in treatment and prophylactic use. Findings obtained by investigating the effects of juglone through more comprehensive *in vivo* studies may change clinicians’ perspectives on treatment.

## Acknowledgments

The authors are grateful to the Afyonkarahisar Health Sciences University Research Foundation, for financial support (Grant number: 21.GENEL.009).

## Data availability statement

The data that support the findings of this study are available from the corresponding author upon reasonable request.

